# Inflammatory monocytes promote pyogranuloma formation to counteract *Yersinia* blockade of host defense

**DOI:** 10.1101/2022.02.06.479204

**Authors:** Daniel Sorobetea, Rina Matsuda, Stefan T. Peterson, James P. Grayczyk, Indira Rao, Elise Krespan, Matthew Lanza, Charles-Antoine Assenmacher, Daniel Beiting, Enrico Radaelli, Igor Brodsky

**Affiliations:** Department of Pathobiology, School of Veterinary Medicine, University of Pennsylvania, Philadelphia, USA

## Abstract

Granulomas are organized immune cell aggregates that form in response to chronic infection or antigen persistence. *Yersinia pseudotuberculosis* (*Yp*) blocks innate inflammatory signaling and phagocytosis, inducing formation of neutrophil-rich pyogranulomas within lymphoid tissues. Here, we uncover that *Yp* triggers pyogranuloma formation within the murine intestinal mucosa, a site not known to contain such structures. Mice lacking circulating monocytes fail to form defined pyogranulomas, have defects in neutrophil activation, and succumb to *Yp* infection. *Yersinia* lacking the virulence factors that block phagocytosis did not induce pyogranulomas, indicating that intestinal pyogranulomas form in response to *Yp* disruption of phagocytosis. Notably, mutation of a single anti-phagocytic virulence factor, YopH, restored pyogranuloma formation and control of *Yp* infection in monocyte-deficient mice, demonstrating that monocytes override YopH-dependent blockade of innate immune defense. This work reveals an unappreciated site of *Yersinia* intestinal invasion, and defines host and pathogen drivers of intestinal granuloma formation.

## Introduction

Microbial pathogens utilize diverse mechanisms to subvert host immunity in order to replicate and spread to new hosts. While acute infections can be cleared rapidly by the immune system, some pathogens evade immune defenses to cause chronic disease. Inability to clear an infection often results in an alternative strategy of containment within structures termed granulomas to limit pathogen dissemination and tissue damage^1^. Granulomas are characterized by the presence of activated phagocytes, notably monocytes and macrophages, and form in response to a wide variety of infectious organisms^2^. Monocytes are rapidly recruited to infected tissues where they produce inflammatory cytokines and anti-microbial effector molecules, contributing to clearance of multiple pathogens^3–8^. Some pathogens, however, exploit monocytes as a means of dissemination, including *Salmonella enterica*, *Yersinia pestis,* and *Mycobacterium* species^9–12^. The pathogen-specific signals that induce granuloma formation remain poorly defined, raising questions about the precise bacterial and host factors that mediate granuloma formation.

The enteropathogenic *Yersinia, Y. pseudotuberculosis* (*Yp*) and *Y. enterocolitica* (*Ye*), cause self-limiting gastroenteritis and mesenteric lymphadenopathy following enteric infection^13, 14^. In immune-compromised patients, bacteria can disseminate systemically to cause a fatal plague-like disease, indicating that the intestinal immune system is critical for control of acute infection. Indeed, studies in mice demonstrate that the intestinal compartment constitutes a major bottleneck against *Yersinia* dissemination, and that bacteria in systemic organs originate predominantly from the intestine rather than gut-associated lymphoid tissues such as the mesenteric lymph nodes (MLN)^15^. A hallmark of *Yersinia* infections is the presence of chronic inflammatory lesions in lymphoid tissue, termed pyogranulomas (PG)^16, 18, 22^. Like classical granulomas, pyogranulomas are characterized by nodular infiltrates of activated phagocytes, including monocytes and macrophages. Pyogranulomas in particular also contain a core of activated neutrophils. How these structures form and which cell types are required for this process are not known. Notably, the contribution of monocytes to pyogranuloma formation and their role in *Yersinia* restriction remain to be fully understood^17, 18^.

Interestingly, pyogranulomas have been observed in the human intestine in response to both *Yp* and *Ye* infection^14, 16, 19–21^ but have not been observed in murine infection models. Here we identify the formation of pyogranulomas in the murine intestinal mucosa during acute enteric *Yersinia* infection. These lesions are enriched in neutrophils and inflammatory monocytes, and contain comparable numbers of live bacteria to Peyer’s patches (PP). Notably, CCR2-deficient mice, which are deficient in circulating inflammatory monocytes^37, 38^, fail to form defined PG, are unable to contain bacteria within the lesions, and succumb rapidly to infection. Moreover, we find that mice lacking CCR2 exhibit reduced levels of IL-1 cytokines and blunted neutrophil activation within intestinal pyogranulomas. Critically, *Yp* lacking either the virulence plasmid itself, which encodes the type III secreted *Yersinia* Outer Proteins (Yops) present in pathogenic *Yersinia*, or lacking the Yops that block phagocytosis, do not induce detectable pyogranulomas, indicating that pyogranulomas are triggered by *Yersinia* blockade of phagocytosis. Moreover, bacterial restriction and formation of defined granulomatous lesions in CCR2-deficient mice is restored in infections with a YopH mutant unable to block phagocytosis, indicating that inflammatory monocytes enable the host immune system to overcome *Yersinia* blockade of phagocytosis. Altogether, our study identifies an unappreciated site of *Yersinia* colonization within the murine intestinal mucosa, and reveals an essential function for inflammatory monocytes in establishing pyogranuloma architecture that enables *Yersinia* clearance.

## Results

Intestinal pyogranulomas form acutely following oral Yersinia infection Yersinia pseudotuberculosis colonizes gut-associated lymphoid tissues, resulting in pyogranuloma formation within five days following oral infection^22^. Interactions between *Yersinia* and immune cells within systemic tissues have been extensively documented^23–25^. In our efforts to investigate the interaction between *Yp* and the intestinal immune system, we observed numerous nodular lesions in the gastrointestinal tract five days post-infection, which appeared as punctate areas of increased opacity along the length of the intestine (Fig. 1a). The total number of lesions in any individual mouse ranged from two to over forty, with a mean of twenty. The lesions were non-uniformly distributed along the intestinal tract and were over-represented in the jejunum and ileum (Extended Data Fig. 1a). Histologic analysis of *Yp-*infected intestines revealed focal regions of inflammation characterized by crypt hyperplasia, edema, and submucosal to transmural cellular infiltration, which contrasted with non-lesional areas of the infected intestine that appeared largely unaffected, similar to uninfected controls (Fig. 1b). Furthermore, the lesions contained infiltrates of macrophages and neutrophils surrounding colonies of coccobacilli, similar to structures we and others have observed in lymphoid tissues^22, 26^, and were therefore termed pyogranulomas (Fig. 1c). Consistently, flow-cytometric analysis of intestinal punch biopsies containing pyogranulomas (PG+), adjacent non-granulomatous tissue (PG-), and uninfected control tissue (uninf) revealed that neutrophils were the most enriched cell type specifically within PG+ tissue, followed by macrophages and inflammatory monocytes (Fig. 1d, Extended Data Fig. 1b). Only minor changes were observed in the frequency or number of eosinophils, dendritic cells, B cells, and T cells in PG+ compared to PG- or uninfected tissue (Extended Data Fig. 1c).

**Figure 1.**
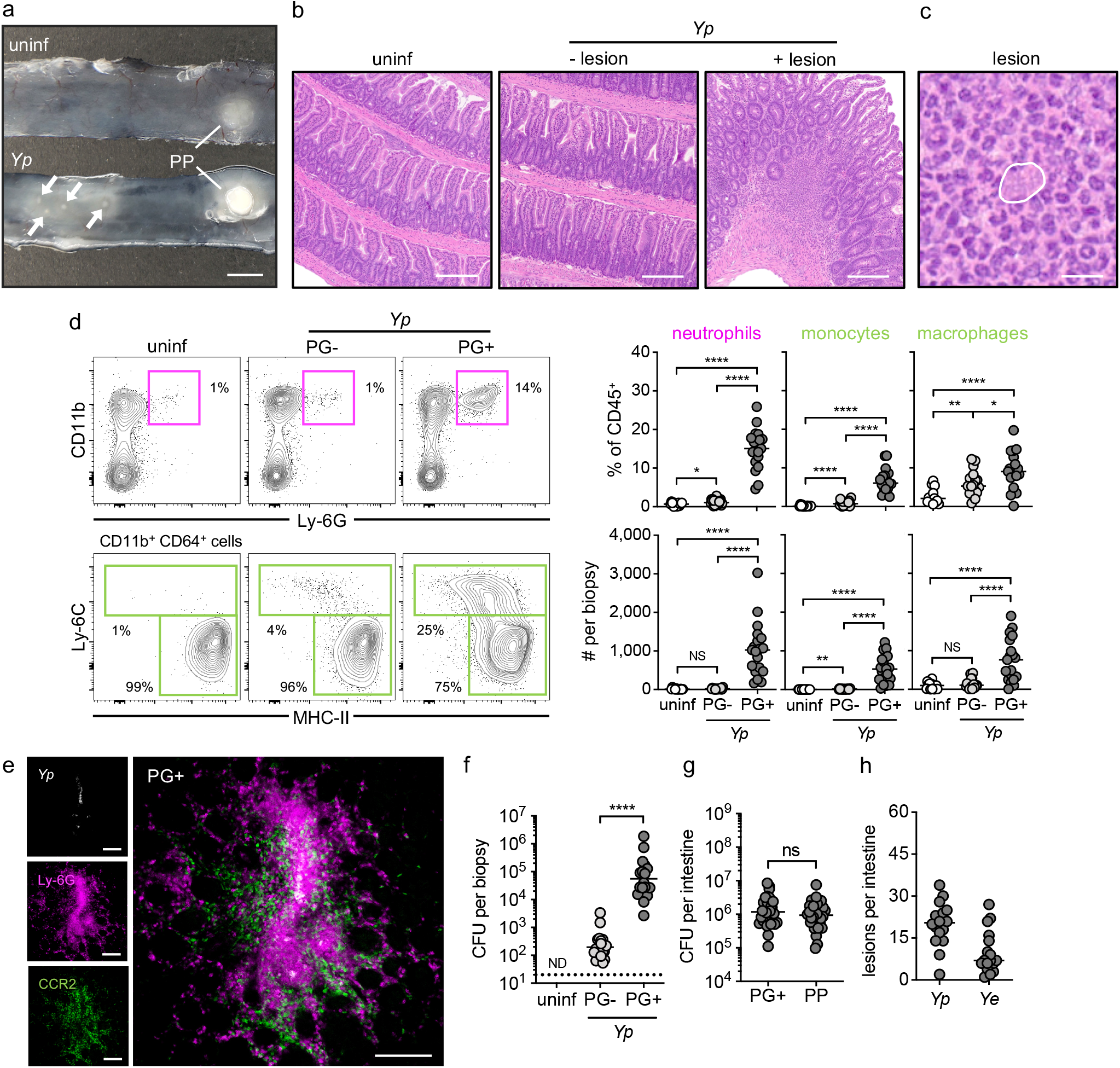
Intestinal pyogranulomas form acutely following oral *Yersinia* infection. (**a**) Longitudinal small-intestinal segments from uninf (top) and *Yp*-infected (bottom) mice with arrows depicting lesions. Scale bar = 3 mm. Representative image of >3 independent experiments. (**b**) H&E-stained paraffin-embedded longitudinal small-intestinal sections from uninf (left) and *Yp*-infected (middle and right) mice. Scale bars = 200 μm. Representative images of three independent experiments. (**c**) H&E-stained paraffin-embedded longitudinal small-intestinal PG+ section from a *Yp*-infected mouse depicting an encircled bacterial colony. Scale bar = 10 μm. Representative image of three independent experiments. (**d**) Flow-cytometry plots (left side) identifying CD11b^+^ Ly-6G^+^ neutrophils (top), CD11b^+^ CD64^+^ Ly-6C^+^ monocytes, and CD11b^+^ CD64^+^ Ly-6C^-^ MHC-II^+^ macrophages (bottom) in small-intestinal uninf (left), PG- (middle) and PG+ (right) tissue with quantification (right side) of frequency (top) and total number (bottom) of neutrophils (left), monocytes (middle) and macrophages (right) in small-intestinal uninf, PG- and PG+ tissue. Each circle represents one mouse. Lines represent median. Pooled data from four independent experiments. Representative images of four independent experiments. (**e**) Fluorescently labeled whole-mount small-intestinal PG+ tissue from an *Yp*- infected *Ccr2^gfp/+^* mouse. White (*Yp-*mCherry), magenta (Ly-6G-AF647), green (CCR2-GFP). Scale bars = 100 μm. Representative images of two independent experiments. (**f**) Bacterial burdens in small-intestinal uninf, PG-, and PG+ tissue. Each circle represents one mouse. Lines represent geometric mean. Pooled data from three independent experiments. (**g**) Cumulative bacterial burdens per intestine in PG+ tissue and PP. Each circle represents one mouse. Lines represent geometric mean. Pooled data from five independent experiments. (**h**) Quantification of total number of intestinal lesions upon *Yp* or *Ye* infection. Each circle represents one mouse. Lines represent median. Pooled data from three independent experiments. Mann-Whitney U test was performed for all statistical analyses. Abbreviations: *Yp* (*Yersinia pseudotuberculosis*), PP (Peyer’s patch), uninf (uninfected control), PG+ (pyogranuloma), PG- (non-granulomatous tissue), H&E (hematoxylin and eosin), CFU (colony-forming units), ND (not detected), NS (not significant, p > 0.05).

*Yp* pyogranulomas in lymphoid tissues consist of a central bacterial colony, surrounded proximally by neutrophils, bordered in turn by monocytes and macrophages^18, 26^. Similarly, confocal microscopy of whole mount PG+ samples demonstrated that intestinal pyogranulomas also contained a central *Yp* colony surrounded by a dense population of Ly-6G^+^ neutrophils (Fig. 1e). Interestingly, in contrast to pyogranuloma architecture in lymphoid tissues, CCR2^+^ monocytes and macrophages formed a dense mesh-like network of cells that both overlapped with and bordered the neutrophils (Fig. 1e). Consistent with the presence of bacterial colonies detected by histology and fluorescence microscopy, PG+ tissue harbored high numbers of viable bacteria, in contrast to neighboring PG- tissue (Fig. 1f). Moreover, the total bacterial burden in PG+ tissue was equivalent to that found in the PP (Fig. 1g). Since Peyer’s patches are viewed as the main entry point and replicative niche for enteropathogenic *Yersinia* following oral innoculation^27, 28^, our data demonstrate that intestinal pyogranulomas represent a previously unappreciated location of *Yersinia* invasion within the intestinal mucosa, and a likely site of bacterial restriction or potential dissemination^29, 30^. Intestinal lesions also formed in response to *Yersinia enterocolitica (Ye)* infection (Fig. 1h), indicating that these lesions are a conserved response to enteric *Yersinia* infection. Together, these findings uncover the first description to our knowledge of intestinal pyogranulomas that contain high numbers of viable bacteria within the murine intestinal mucosa following oral infection and may serve as an intestinal niche for enteric *Yersinia*.

### The intestinal innate inflammatory response to Yersinia is localized to pyogranulomas

Intestinal pyogranulomas represent an unrecognized site for enteric *Yersinia* to interact with the murine mucosal immune system. To define the PG- specific inflammatory response, we performed RNA sequencing of punch biopsies containing pyogranulomas (PG+), adjacent non-pyogranuloma tissue (PG-), and uninfected control tissue. Principal-component analysis showed distinct clustering by sample type (Fig. 2a), and comparison of PG+ and PG- samples revealed 355 upregulated and 363 downregulated genes (Fig. 2b). Top upregulated genes included granulocyte- and monocyte-recruiting chemokines (*Cxcl1, Cxcl2, Cxcl3, Cxcl5*), pro-inflammatory cytokines (*Il1b, Il22*), antimicrobial metal-sequestration proteins (*S100a8, S100a9*), and matrix metalloproteases (*Mmp3*) (Fig. 2c). Gene ontology analysis revealed that chemotaxis of myeloid cells and defense against bacterial pathogens predominated the top upregulated responses within PG+ biopsies (Fig. 2d, Extended Data Table 1). These features of the transcriptional response were strikingly similar to those previously reported in *Yp*-infected PP^31^. Indeed, gene set-enrichment analysis indicated that the reported top 50 upregulated genes in *Yp*-infected PP were also enriched in PG+ samples (Fig. 2e). Consistently, protein levels of pro-inflammatory cytokines TNF, CCL2, IL-1β, IL-1α, and IL-6 were elevated within PG+ biopsies compared to PG- and uninfected biopsies (Fig. 2f). Consistent with the histology and microscopy analyses, the inflammatory transcriptional response was localized to PG+ tissue, as PG- samples did not exhibit enrichment of genes or ontology terms related to myeloid cell migration or innate immune-cell activation (Extended Data Fig. 2, Extended Data Table 2). Likewise, production of pro-inflammatory cytokines was not detected above uninfected levels in PG- tissue (Fig. 2e). Altogether, our transcriptional and cytokine analyses indicate that the pro-inflammatory response to *Yp* infection in the gut mucosa is driven by innate immune cells and spatially restricted to pyogranulomas.

**Figure 2.**
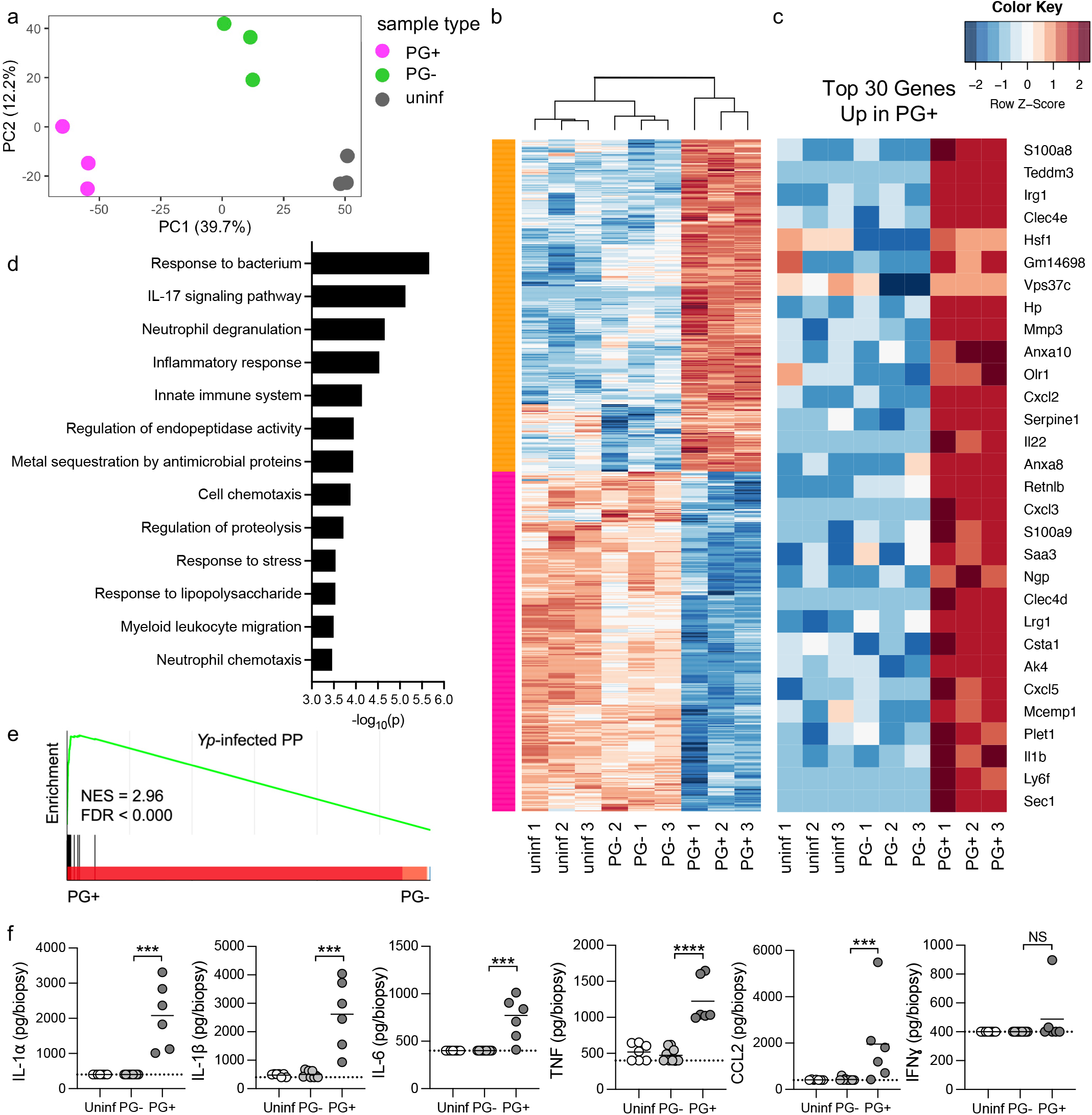
The intestinal innate inflammatory response to *Yersinia* is localized to pyogranulomas. (**a**) Principal component analysis of PG+ (pink), PG- (green), and uninf (gray) samples at day 5 post-infection. Five biopsies were pooled per mouse. (**b**) Heatmap of all differentially expressed genes in PG+ compared to PG- samples. False discovery rate < 0.05 using Benjamini-Hochberg procedure. (**c**) Heatmap of top 30 significantly upregulated genes in PG+ compared to PG- samples in descending order by fold change. False discovery rate < 0.05 using Benjamini-Hochberg procedure. (**d**) Gene ontology analysis of top 30 upregulated genes by fold change only in PG+ compared to PG- samples. (**e**) Gene set enrichment analysis (GSEA) of top 50 upregulated genes in *Yp*-infected Peyer’s patches. NES = normalized enrichment score; FDR = false discovery rate. (**f**) Cytokine levels in homogenates of tissue punch biopsies were measured by cytometric bead array. Lines represent group mean. Mann-Whitney U test was performed for statistical analysis. NS = not significant. Data from one or two pooled independent experiments.

### Inflammatory monocytes are required for organized pyogranuloma formation and restriction of Yersinia within intestinal tissue

Inflammatory monocytes promote host defense by differentiating into phagocytes and antigen presenting cells^3, 5, 7, 32, 33^, producing pro-inflammatory cytokines and antimicrobial molecules^4^, and modulating the function of other immune cells to promote pathogen clearance or prevent immunopathology^6, 34^. However, monocyte-derived cells can also promote pathogen replication or pathogen dissemination to new infection sites^9, 29, 30, 35, 36^. Mice lacking the chemokine receptor CCR2 have a ten-fold reduction in circulating monocytes due to defective bone marrow-egress^37, 38^. Monocytes are rapidly recruited to intestinal pyogranulomas (Fig. 1) and systemic *Yp* infection sites^18, 22, 26, 39, 40^. CCR2-deficiency was reported to be associated with more rapid bacterial clearance from the MLN following enteric infection^18^, but increased susceptibility to intravenous *Yp* infection^17^. We therefore sought to dissect the contribution of inflammatory monocytes to anti-*Yersinia* innate immune defense and formation of pyogranulomas. Although infection of CCR2-deficient mice^41, 42^ resulted in similar numbers of macroscopic intestinal lesions to WT mice (Fig. 3a, Extended Data Fig. 3a), blinded analysis by a board-certified pathologist revealed that intestinal lesions in *Ccr2^gfp/gfp^* mice (in which insertion of EGFP into the translation initiation site of *Ccr2* disrupts its expression) had a disorganized appearance, and many showed a central region of caseation containing substantial tissue necrosis that was not observed in intestinal PG of WT mice (Fig. 3b). Whereas WT pyogranulomas exhibited robust inflammatory infiltrates and a defined cellular organization encapsulating central bacterial colonies, *Ccr2^gfp/gfp^* intestinal lesions contained extensively distributed bacterial colonies with limited inflammatory immune cell recruitment (Fig. 3c). Two independent lines of CCR2-deficient mice had significantly higher bacterial burdens in both PG+ and PG- intestinal tissue biopsies than corresponding tissues of WT mice, suggesting that monocytes restrict enteric *Yp* infection within pyogranulomas and limit dissemination to surrounding intestinal tissue (Fig. 3d, Extended Data Fig. 3b). Consistent with their reduced cellularity in histologic analyses, flow cytometry of *Ccr2^gfp/gfp^* lesions revealed an overall decrease in viable CD45^+^ hematopoietic cells compared to WT pyogranulomas (Fig. 3e). Consistent with the known role of CCR2 in egress of monocytes out of bone marrow and into circulation^37, 38^, *Ccr2^gfp/gfp^* intestinal lesions contained significantly lower numbers of monocytes and macrophages compared with pyogranulomas from WT mice. Surprisingly, despite similar frequencies of neutrophils in the MLN and spleen upon infection (Extended Data Fig. 3c), *Ccr2^gfp/gfp^* intestinal lesions showed reduced neutrophil numbers (Fig. 3e), suggesting that monocytes or monocyte-derived cells promote recruitment, retention, or survival of neutrophils within intestinal pyogranulomas during *Yp* infection. Notably, immunofluorescence microscopy of the lesions revealed that the bacteria expanded outside range of the neutrophil marker Ly-6G, suggesting that neutrophils were unable to effectively contain *Yp* in the absence of monocytes. In contrast, *Yp* was fully encapsulated by neutrophils and CCR2^+^ cells in CCR2-sufficient PGs (Fig. 3f). Intriguingly, neutrophil surface expression of the integrin CD11b, an established marker of neutrophil activation^43–46^, was also significantly reduced in both *Ccr2^gfp/gfp^* PGs and MLN compared to WT mice (Fig. 3g, Extended Data Fig. 3d), indicating a defect in neutrophil activation in these tissues in the absence of monocytes. Moreover, IL-1α and IL-1β levels were significantly decreased in *Ccr2^gfp/gfp^* PGs (Fig. 3h), suggesting that monocytes or monocyte-derived cells were required to either directly release IL-1 or promote IL-1 production by other cells within intestinal PGs. CCR2-deficiency specifically impacted IL-1 levels within PGs, as levels of other pro-inflammatory cytokines was not altered in CCR2-deficient mice (Extended Data Fig. 3e). Importantly, we found no effect of CCR2 deficiency on CD4^+^ or CD8^+^ T cell numbers^47^ in *Ccr2^gfp/gfp^* intestinal PGs, suggesting that the defect in enteric control of *Yp* in CCR2-defficient mice was not due an effect on T cell numbers (Extended Data Fig. 3f). Altogether, these results demonstrate that monocyte-derived cells are required to establish robust, functional granulomas that limit intestinal bacterial replication and dissemination.

**Figure 3.**
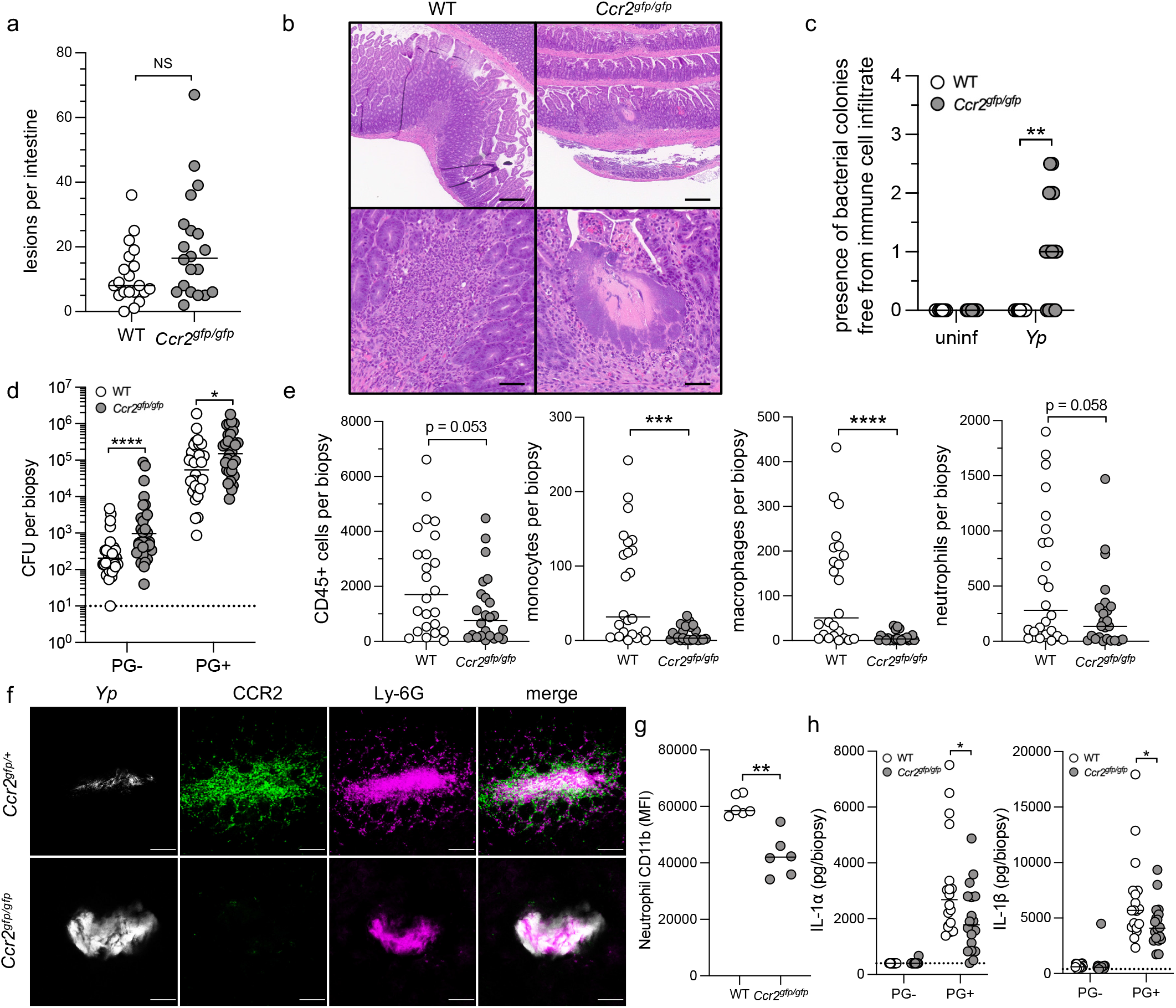
Inflammatory monocytes are required for organized pyogranuloma formation and restriction of Yersinia within intestinal tissue. (**a**) Quantification of total number of intestinal lesions at day 5 post infection. Each circle represents one mouse. Lines represent median. Pooled data from four independent experiments. (**b**) H&E-stained paraffin-embedded longitudinal small intestinal sections from *Yp*- infected mice. Scale bars = 250 μm (top) and 50 μm (bottom). Representative images of two independent experiments. (**c**) Histopathological scores of small intestinal tissue from uninfected and *Yp*- infected mice. Each mouse was given a score between 0-4 (healthy-severe) for signs of pathology. Each circle represents one mouse. Lines represent median. Pooled data from two independent experiments. (**d**) Bacterial burdens in small-intestinal PG- and PG+ tissue. Each circle represents the mean *Yp*-CFU of 3-5 pooled punch biopsies from one mouse. Lines represent geometric mean. Pooled data from six independent experiments. (**e**) Total number of CD45^+^ cells, monocytes, macrophages, and neutrophils in small intestinal uninf, PG-, and PG+ tissue. Each circle represents the mean of 3-10 pooled punch biopsies from one mouse. Lines represent median. Pooled data from five independent experiments. (**f**) Fluorescently labeled whole-mount small intestinal PG+ tissue from *Yp*-infected *Ccr2^gfp/+^* (top) and *Ccr2^gfp/gfp^* (bottom) mice. Scale bars = 100 μm. Representative images of two independent experiments. (**g**) PG+ neutrophil CD11b expression was measured by flow cytometry. Lines represent median. Representative of four independent experiments. (**h**) Cytokine levels in homogenates of tissue punch biopsies were measured by cytometric bead array. Lines represent median. Pooled data from three independent experiments. Mann-Whitney U test was performed for all statistical analyses. NS = not significant.

### Inflammatory monocytes are required for systemic control of Yersinia and host survival

Following dissemination from the intestine, *Yp* colonizes and induces PG formation in lymphoid tissues^18, 22, 26^. Critically, CCR2-deficient mice had significantly higher *Yp* burdens in the MLN and systemic organs (Fig. 4a, Extended Data Fig. 3g). Histopathological analysis revealed that similar to intestinal tissues, infected *Ccr2^gfp/gfp^* spleens lacked defined pyogranulomas and exhibited widespread tissue necrosis, free bacterial colonies, and sparse immune cell recruitment, which contrasted with WT spleens that contained neutrophils and monocytes that effectively surrounded *Yp* microcolonies (Fig. 4b-c). Notably, mice lacking CCR2 succumbed rapidly to acute infection (Fig. 4d, Extended Data Fig. 3h). Importantly, infection of co-housed littermate progeny from heterozygous mating demonstrated that *Ccr2^+/+^* and *Ccr2^gfp/+^* mice were equally resistant to *Yp* infection while *Ccr2^gfp/gfp^* littermates succumbed (Fig. 4e), indicating that differences in intestinal microbiota between *Ccr2^gfp/gfp^* and WT mice were unlikely to account for differences in susceptibility to *Yp* infection. Collectively, our findings demonstrate that inflammatory monocytes are required for formation of organized pyogranulomas in host tissues, thereby limiting tissue necrosis and systemic bacterial dissemination, ultimately enabling bacterial control and host survival following oral *Yp* infection.

**Figure 4.**
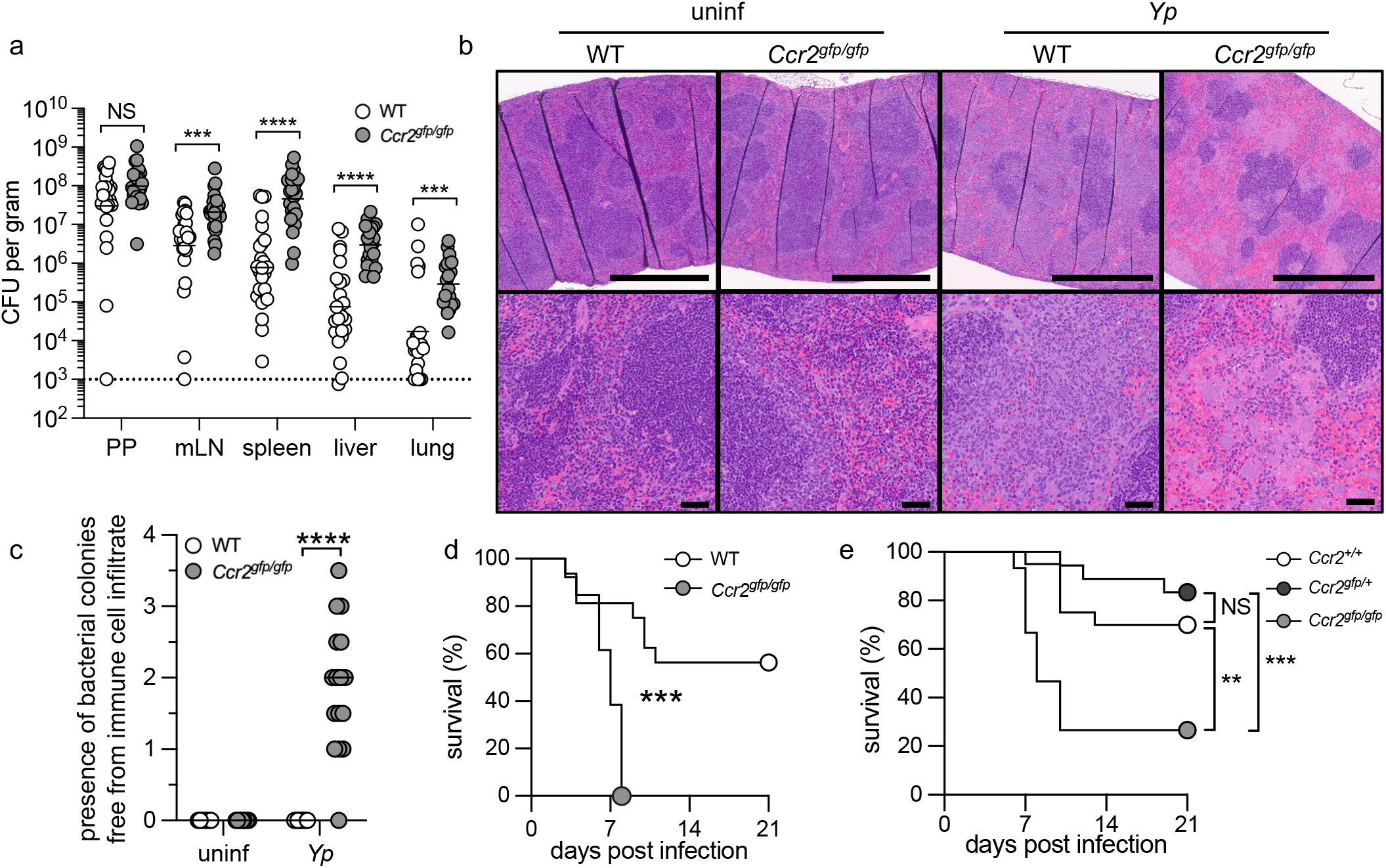
Inflammatory monocytes are required for systemic control of *Yersinia* and host survival. (**a**) Bacterial burdens in indicated organs at day 5 post infection. Each circle represents one mouse. Lines represent geometric mean. Pooled data from four independent experiments. (**b**) H&E-stained paraffin-embedded longitudinal spleen sections from WT and *Ccr2^gfp/gfp^* mice. Scale bars = 500 μm (top) and 50 μm (top). Representative images of two independent experiments. (**c**) Histopathological scores of spleens from uninfected and *Yp*-infected mice. Each mouse was given a score between 0-4 (healthy-severe) for signs of pathology. Each circle represents one mouse. Lines represent median. Pooled data from two independent experiments. (**d**) Survival of infected WT (n=16) and *Ccr2^gfp/gfp^* (n=13) mice. Pooled data from two independent experiments. (**e**) Survival of infected littermate wild-type *Ccr2^+/+^* (n=20), heterozygous *Ccr2^gfp/+^* (n=19), and homozygous *Ccr2^gfp/gfp^* (n=15) mice. Pooled data from three independent experiments. Statistical analyses by (a, c) Mann-Whitney U test and (d-e) Mantel-Cox test. NS = not significant.

### Intestinal pyogranulomas form in response to Yersinia virulence

Mammalian pathogenic *Yersinia* inject their target cells with effector proteins (Yops) that are encoded on a virulence plasmid (pYV)^48^. Yops block antimicrobial defenses, including phagocytosis, production of reactive oxygen species (ROS), and degranulation^24, 49–52^. *Yp* lacking the virulence plasmid (pYV-) are still able to colonize gut-associated lymphoid tissues, including the MLN, during the acute phase of infection without causing disease^53^. As granuloma formation occurs in response to pathogens that thwart immune defenses, we hypothesized that formation of intestinal pyogranulomas is triggered by the activity of *Yp* effector proteins. Indeed, even at a ten-fold higher infectious dose, we were unable to detect any macroscopic intestinal lesions at day 5 post-infection with pYV-bacteria (Fig. 5a), despite significant, albeit lower, bacterial burdens in the intestinal mucosa (Fig. 5b). We likewise found lower bacterial burdens within gut-associated and systemic lymphoid tissues of mice infected with pYV-compared to WT bacteria (Fig. 5c). Intriguingly, monocytes were still recruited to both the intestinal mucosa and MLN in response to of pYV-infection, indicating that absence of detectable pyogranulomas was not due to an absolute lack of inflammatory cell infiltration into the tissue of pYV-infected mice (Fig. 5d,e). Notably, however, neutrophil recruitment to the intestine or MLN was absent in response to pYV-infection, demonstrating that neutrophil recruitment or retention in the intestinal tissue is a response to *Yp* virulence activities (Fig. 5d,e).

**Figure 5.**
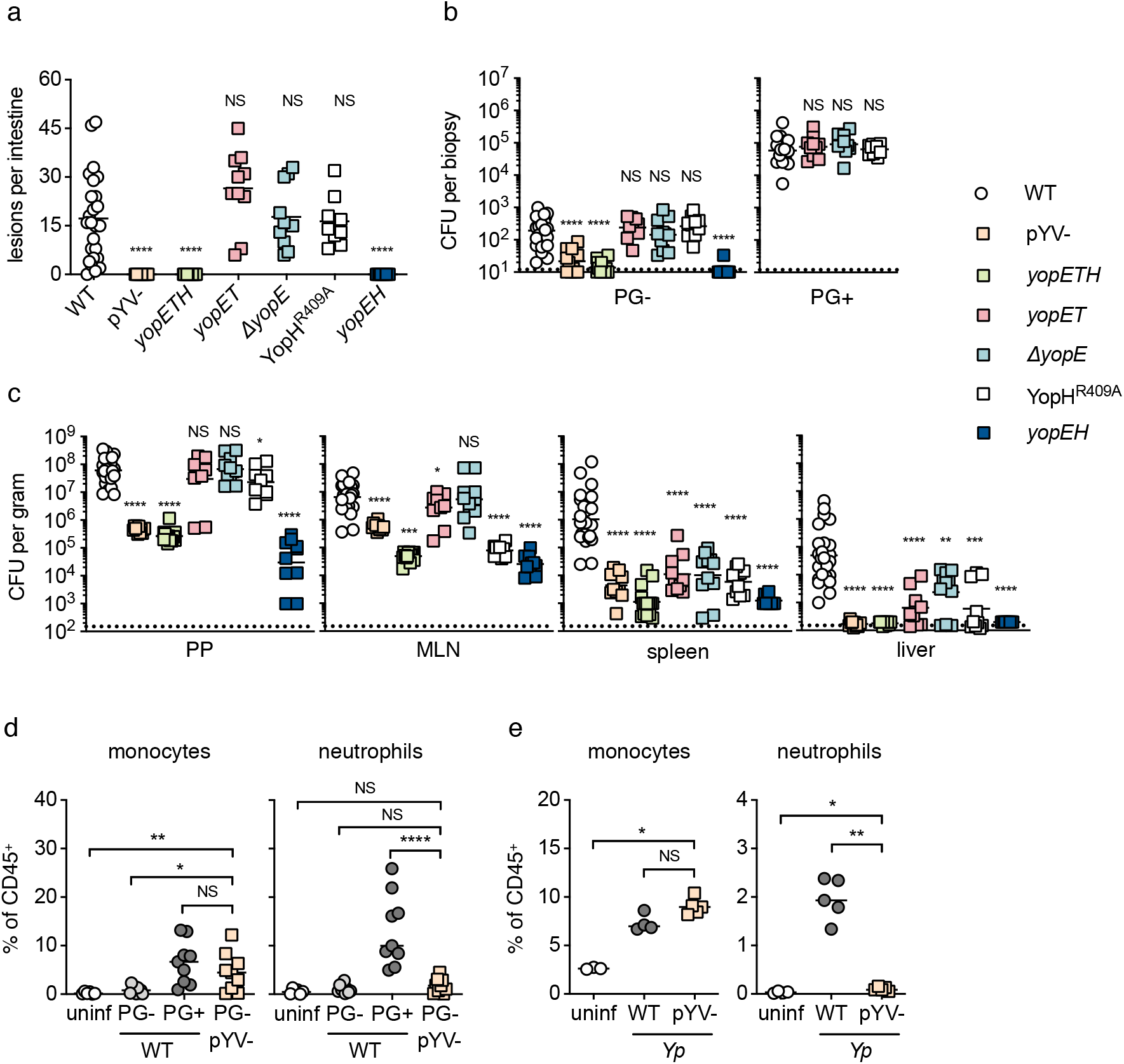
Intestinal pyogranulomas form in response to *Yersinia* virulence. (**a**) Quantification of total number of intestinal lesions upon infection. Each symbol represents one mouse. Lines represent median. Pooled data from 2-3 independent experiments. (**b**) Bacterial burdens in small-intestinal PG- (left) and PG+ (right) tissue. Each symbol represents one mouse. Lines represent geometric mean. Pooled data from 2-3 independent experiments. (**c**) Bacterial burdens in indicated organs. Each symbol represents one mouse. Lines represent geometric mean. Pooled data from 2-3 independent experiments. (**d**) Frequency of monocytes (left) and neutrophils (right) in small-intestinal uninf, PG- and PG+ tissue. Each symbol represents one mouse. Lines represent median. Pooled data from two independent experiments. (**e**) Frequency of monocytes (left) and neutrophils (right) in MLN. Each symbol represents one mouse. Lines represent median. Representative data from two independent experiments. Mann-Whitney U test was performed for all statistical analyses.

To test whether specific effectors were responsible for triggering pyogranuloma formation, we infected mice with individual or combined mutants of different Yops. Intestinal lesions formed at similar numbers in mice infected with *Yp* lacking either YopM (*ΔyopM*) or deficient in YopJ enzymatic activity (YopJ^C172A^), which block pyrin inflammasome assembly and MAPK signaling^54–60^, respectively, suggesting that neither YopM nor YopJ are solely required to trigger pyogranuloma formation (Extended Data Fig. 4a). A critical virulence function contained on the pYV plasmid is disruption of the actin cytoskeleton, leading to blockade of phagocytosis and inhibition of the reactive oxygen burst^24, 61–63^. These Yops (E, H, and T), have partially overlapping functions and can compensate for one another in certain settings^62–69^. Notably, *Yp* containing point mutations in each of the key catalytic residues of these Yops (YopE^R144A^, YopT^C139A^, and YopH^R409A^) did not induce intestinal lesions and had burdens similar to pYV-infection (Fig. 5a-c). In contrast, bacteria lacking only YopE (*ΔyopE*) or both YopE and YopT (*yopET*) induced formation of similar numbers of intestinal lesions (Fig. 5a) and had wild-type levels of intestinal colonization (Fig. 5b,c). These data indicate that YopH is sufficient, in the absence of YopE and YopT, to induce pyogranuloma formation. Interestingly, bacteria lacking only YopH enzymatic activity or YopE alone had similar numbers of macroscopic lesions to wild-type-infected mice (Fig. 5a), suggesting that YopH is sufficient, but not absolutely required to trigger pyogranuloma formation. Notably, while bacteria lacking either YopE or YopH alone induced similar numbers of intestinal lesions to wild-type *Yersinia*, bacteria lacking both YopE and YopH (*yopEH*) induced no detectable lesions, and showed similar levels of colonization to *yopETH* bacteria (Fig. 5a-c). Altogether, these data indicate that intestinal pyogranuloma formation occurs in response to *Yersinia* disruption of host actin cytoskeletal dynamics and is primarily driven by YopE and YopH.

### Yp restriction by CCR2-deficient mice is restored in absence of YopH activity

Our results demonstrate that YopH is sufficient, but not strictly necessary to induce formation of intestinal pyogranulomas. YopH potently blocks phagocytosis by innate immune cells, including macrophages and neutrophils^24, 49, 50, 62, 69, 70^. YopH-deficient *Yp* therefore have enhanced susceptibility to neutrophil killing *in vitro* and are attenuated *in vivo*^71–73^. While YopH potently blocks neutrophil functions, neutrophil depletion increases susceptibility to *Yp* and restores virulence to *yopH-*mutant bacteria^50, 70^, indicating that neutrophil defenses nevertheless counteract YopH activity *in vivo*. How neutrophils counteract the activity of YopH is not understood. Monocyte and neutrophil crosstalk is essential for optimal immune defense in many infectious contexts^74^. However, whether monocytes are required to combat YopH-dependent blockade of neutrophil function is unknown. To test this possibility, we infected CCR2-deficient mice with WT or YopH-mutant *Yp*. Strikingly, CCR2-deficient mice infected with YopH^R409A^ bacteria exhibited robust neutrophil recruitment and containment of bacterial colonies similar to WT mice, in contrast to CCR2-deficient mice infected with WT *Yp* (Fig. 6a). Moreover, both *Ccr2^gfp/gfp^* and *Ccr2^-/-^* mice restricted YopH^R409A^ bacteria within the intestine (Fig. 6b, Extended Data Fig. 4a) and systemically (Fig. 6c, Extended Data Fig. 4b). While CCR2-deficient mice rapidly succumbed to WT *Yp*, they survived infection with YopH^R409A^ bacteria similarly to WT mice (Extended Data Fig. 4c). Altogether, our results uncover a key role for monocytes in establishment and function of intestinal pyogranulomas, revealing monocytes as essential in overcoming *Yersinia* suppression of neutrophil function *in vivo* and preventing acute mortality.

**Figure 6.**
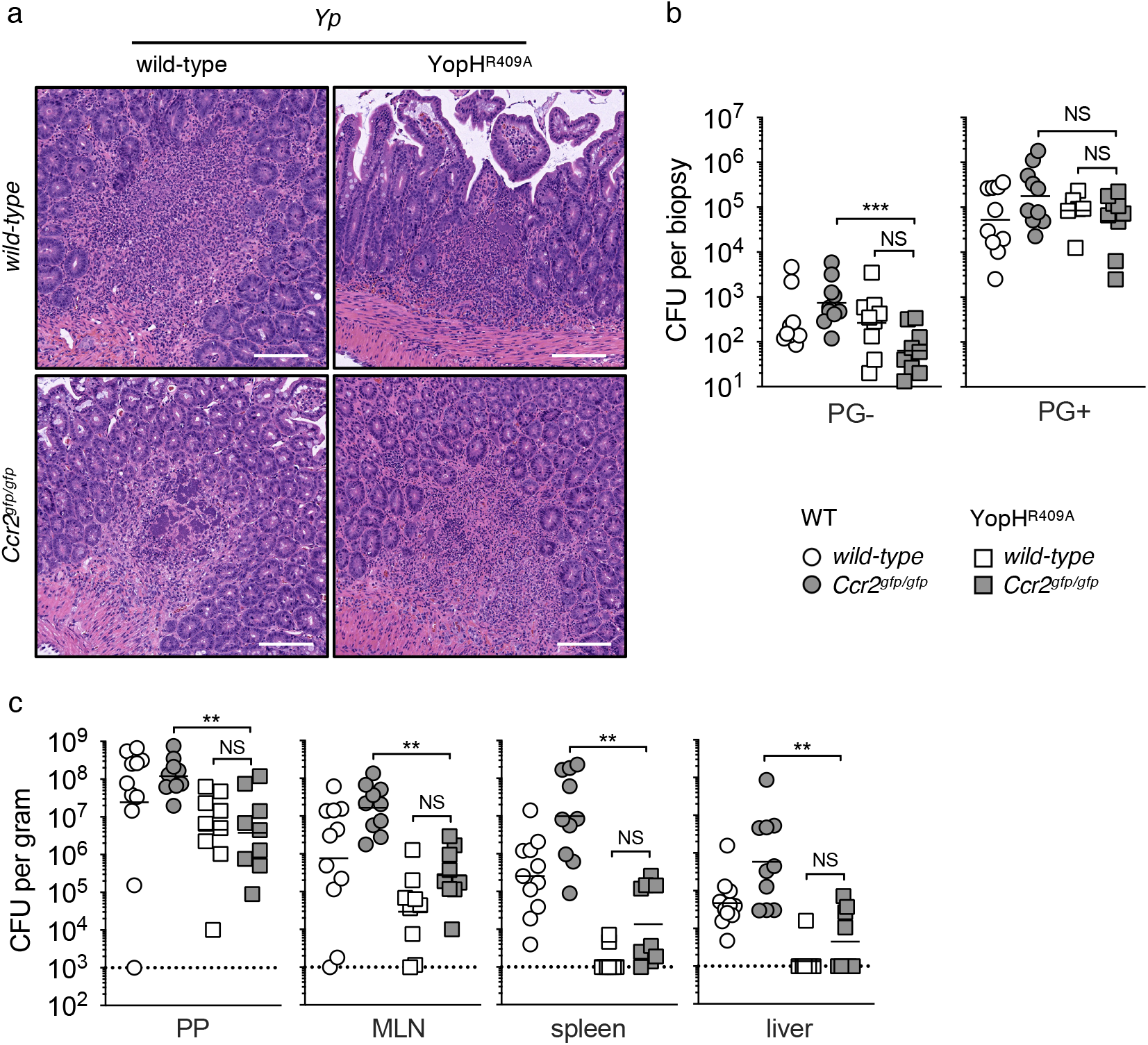
*Yp* restriction by CCR2-deficient mice is restored in absence of YopH activity. (**a**) H&E-stained paraffin-embedded transverse small intestinal sections from WT (top) and *Ccr2^gfp/gfp^* (bottom) mice infected with WT (left) or YopH^R409A^ (right) *Yp* at day 5 post infection. Scale bars = 100 μm. Representative images of two independent experiments. (**b**) Bacterial burdens in small-intestinal PG- and PG+ tissue at day 5 post infection. Each symbol represents one mouse. Lines represent geometric mean. Pooled data from two independent experiments. (**c**) Bacterial burdens in indicated organs at day 5 post infection. Each symbol represents one mouse. Lines represent geometric mean. Pooled data from two independent experiments. Mann-Whitney U test was performed for all statistical analyses. NS = not significant.

## Discussion

Granulomas are a conserved immunological response to persistent or long-term infectious and non-infectious stimuli^1, 2^. The natural rodent and human pathogen *Yersinia pseudotuberculosis* induces formation of neutrophil-rich lesions termed pyogranulomas in infected lymphoid tissues^14, 16^. Here, we report that pyogranulomas form throughout the intestine during murine enteropathogenic *Yersinia* infection. Peyer’s patches are considered the primary site of intestinal infection by *Yersinia*. However, intestinal pyogranulomas harbored a similar total bacterial burden as the PPs, suggesting that PGs are a previously unrecognized niche for enteropathogenic *Yersinia* intestinal colonization. Notably, wild-type mice almost entirely restricted intestinal *Yp* and inflammation to these granulomatous foci. The pro-inflammatory transcriptional profile of PGs was similar to *Yp-*infected Peyer’s patches^31^, suggesting a shared response to intestinal *Yp* infection driven by the recruitment and activation of innate immune cells. These structures may therefore serve to limit bacterial spread and tissue damage within the gut mucosa.

We found that monocyte-derived cells were critical for the architectural integrity of PGs in the intestine and deeper tissues during *Yersinia* infection. While other cell types express CCR2, notably activated T cells, we did not observe altered numbers of these cells within intestinal PGs in CCR2-deficient mice. Absence of circulating monocytes was associated with widespread tissue necrosis and elevated bacterial burdens in intestinal and systemic tissues, indicating that the monocyte lineage is critical for containment of *Yersinia* infection. Furthermore, CCR2-deficiency was associated with defective pro-inflammatory IL-1 production and neutrophil activation within intestinal PGs.

A previous study reported that monocytes were dispensable for acute control of *Yp* following oral inoculation^18^. These studies employed a distinct *Ccr2^-/-^* mouse line^32, 38^, originally generated on the 129 mouse background and backcrossed to C57BL/6^37^, raising the possibility that distinct polymorphisms in this mouse line might account for this difference. *Ccr2^gfp/gfp^* mice were generated on the C57BL/6J background^41^, making it unlikely that our findings are due to immunologically impactful polymorphisms that co-segregate with the *Ccr2* locus. Infection of co-housed littermates also demonstrated that increased susceptibility of CCR2-deficient mice is not due to differences in maternally-transmitted or environmentally-acquired intestinal microbiota.

Formation of intestinal PGs also required the anti-phagocytic Yops. While combined deficiency in two effector proteins that block phagocytosis, Yops E and H, abrogated PG formation, either effector alone was sufficient to induce PG formation. This suggests that PG formation occurs in response to pathogens that resist immune clearance, and that the anti-phagocytic activities of YopH and YopE independently mediate PG formation. Precisely how blockade of phagocytosis by YopsH and E lead to recruitment of phagocytic cells and assembly of pyogranulomas remains to be determined in future studies.

Monocytes and neutrophils comprise a large proportion of Yop-injected cells *in vivo*^75^. YopH blocks neutrophil degranulation, ROS production, phagocytosis, and release of neutrophil extracellular traps^24, 50, 70, 73, 76^. Notably, while monocytes were required for host survival in response to WT bacteria, they were dispensable for control of *Yp* lacking YopH activity. Our findings imply that monocytes are critical for host defense against pathogen blockade of neutrophil function, either by directly restricting bacterial growth or enhancing neutrophil-mediated bacterial control. Indeed, intestinal PGs in CCR2-deficient mice exhibited decreased neutrophil numbers and activation, suggesting that monocytes promote neutrophil function during intestinal *Yp* infection. Altogether, our study reveals an unappreciated site of *Yersinia*-host interaction within the intestine, and provides insight into mechanisms of *Yersinia* granuloma formation and function.

## Methods

### Animals

C57BL/6 wild-type and *Ccr2^gfp/gfp^* mice^41^ (in which insertion of EGFP into the translation initiation site of *Ccr2* disrupts its expression) were acquired from the Jackson Laboratory and bred at the University of Pennsylvania. *Ccr2*^-/-^ mice^42^ were provided by Dr. Sunny Shin (University of Pennsylvania). Unless specifically noted, all animals were bred by homozygous mating and housed separately by genotype. All animal studies were performed in strict accordance with University of Pennsylvania Institutional Animal Care and Use Committee-approved protocols.

### Bacteria

Wild-type *Yp* (strain 32777, serogroup O1)^77^ and isogenic mutants^58, 77–79^ were provided by Dr. James Bliska (Dartmouth College). *Ye* (strain 8081, serogroup O8)^80^ was provided by Dr. Stanley Falkow (Stanford University). Additional mutants lacking YopE (*ΔyopE*), enzymatic activity of YopH (YopH^R409A^) or YopE/YopT/YopH (YopE^R144A^T^C139A^H^R409A^, denoted *yopETH*) were generated by two-step allelic recombination as previously described^79^ with plasmids provided by Dr. James Bliska. Fluorescent *Yp* (mCherry^+^) was generated from plasmids provided by Dr. Kimberly Davis (Johns Hopkins University).

### Infections

*Yp* and *Ye* were cultured to stationary phase at 28°C and 250 rpm shaking for 16 hours in 2xYT broth supplemented with 2 μg/ml triclosan (Millipore Sigma). Mice of either sex between 8-12 weeks of age were fasted for 16 hours and subsequently inoculated by oral gavage with 100-200 μl phosphate-buffered saline (PBS). All bacterial strains were administered at 2×10^8^ colony-forming units (CFU) per mouse with the exception of pYV-which was administered at 20×10^8^ CFU per mouse.

### Bacterial CFU quantifications

In order to determine bacterial burdens, tissue biopsies were collected in sterile PBS, weighed, homogenized for 40 seconds with 6.35 mm ceramic spheres (MP Biomedical) using a FastPrep-24 bead beater (MP Biomedical) and serially diluted tenfold in PBS before being plated on LB agar supplemented with 2 μg/ml triclosan and incubated for two days at room temperature. Dilutions of each sample were plated in triplicate and expressed as the mean CFU per gram of tissue or per biopsy.

### Protein quantifications

Cytokines were measured in supernatants from homogenized tissue using Cytometric Bead Array (BD Biosciences) according to manufacturer’s instructions with the following modification: the amount of capture beads, detection reagents, and sample volumes was scaled down tenfold. Data were collected on an LSRFortessa flow cytometer (BD Biosciences) and analyzed with FlowJo v10 (BD Biosciences).

### Tissue preparation and cell isolation

Lymph nodes were homogenized in R10 buffer consisting of RPMI 1640 (Millipore Sigma) supplemented with 10 mM HEPES (Millipore Sigma), 10% fetal bovine serum (Omega Scientific), 1 mM sodium pyruvate (Thermo-Fisher Scientific) and 100 U/ml penicillin + 100 μg/ml streptomycin (Thermo Fisher Scientific), then passed through 70 μm cell strainers (Fisher Scientific). Intestines were excised, flushed luminally with sterile PBS to remove the feces, opened longitudinally along the mesenteric side and placed luminal side down on cutting boards (Epicurean). Small-intestinal tissue containing pyogranulomas (PG+), adjacent non-granulomatous areas (PG-) and uninfected control tissue (uninf) were excised using a 2 mm-ø dermal punch-biopsy tool (Keyes). Biopsies within each mouse were pooled groupwise, suspended in epithelial-dissociation buffer consisting of calcium and magnesium-free HBSS (Thermo Fisher Scientific) supplemented with 15 mM HEPES, 10 mg/ml bovine serum albumin (Millipore Sigma), 5 mM EDTA and 100 U/ml penicillin + 100 μg/ml streptomycin, then incubated for 30 minutes at 37°C under continuous agitation. To isolate immune cells from the lamina propria, the tissue was enzymatically digested in R10 buffer, along with 0.5 Wünsch units/ml liberase TM (Roche), 30 μg/ml DNase I (Roche), and 5 mM CaCl_2_ for 20 min at 37°C under continuous agitation. The resulting cell suspensions were filtered through 100 μm cell strainers (Fisher Scientific) and subjected to density-gradient centrifugation using Percoll (GE Healthcare). Briefly, cells were suspended in 40% Percoll and centrifuged over a 70% Percoll layer for 20 min at 600 × g with lowest brake at room temperature. Cells collected between the layers were washed with R10 for downstream analysis.

### Flow cytometry

Unspecific Fc binding was blocked for 15 minutes on ice with unconjugated anti-CD16/CD32 (clone 93; Thermo-Fisher Scientific). Cells were subsequently fluorescently labeled for 30 minutes on ice with the following antibodies and reagents: PE-conjugated rat anti-mouse Siglec-F (E50-2440; BD Biosciences), PE-TxR-conjugated rat anti-mouse CD11b (M1/70.15; Thermo Fisher Scientific), PE-Cy5-conjugated mouse anti-mouse NK1.1 (PK136; BioLegend), PE-Cy5.5-conjugated rat anti-mouse CD4 (RM4-5; Thermo Fisher Scientific), PE-Cy7-conjugated rat anti-mouse CD3 (17A2; BioLegend), FITC-conjugated Armenian hamster anti-mouse CD11c (N418; BioLegend), PerCP-Cy5.5-conjugated rat anti-mouse Ly-6C (HK1.4; Thermo Fisher Scientific), PB-conjugated rat anti-mouse CD90.2 (53-2.1; BioLegend), BV510-conjugated rat anti-mouse CD19 (1D3; BD Biosciences), BV605-conjugated Armenian hamster anti-mouse TCRβ (H57-597; BD Biosciences), BV650-conjugated rat anti-mouse I-A/I-E (M5/114.15.2; BD Biosciences), BV711-conjugated rat anti-mouse CD8α (53-6.7; BD Biosciences), BV785-conjugated rat anti-mouse Ly-6G (1A8; Thermo Fisher Scientific), AF647-conjugated mouse anti-mouse CD64 (X54-5/7.1; BD Biosciences) and AF700-conjugated mouse anti-mouse CD45.2 (104; BioLegend) along with eF780 viability dye (BioLegend) diluted in PBS according to manufacturer’s instructions. Cells were fixed for 20 minutes on ice with Cytofix/Cytoperm Fixation/Permeabilization solution (BD Biosciences). Cells were acquired on an LSRFortessa flow cytometer and data was analyzed with FlowJo v10. Dead and clustered cells were removed from all analyses.

### Histology

Tissues were fixed in 10% neutral-buffered formalin (Fisher Scientific) and stored at 4°C until further processed. Tissue pieces were embedded in paraffin, sectioned by standard histological techniques and stained with hematoxylin and eosin for subsequent histopathological disease scoring by blinded board-certified pathologists. Tissue sections were given scores between 0-4 (healthy-severe) for multiple parameters, including degree of inflammatory cell infiltration, necrosis, and free bacterial colonies along with tissue-specific parameters such as villus blunting and crypt hyperplasia. Healthy mice were characterized by and subsequently scored as having none or low levels of the parameters described, whereas severely afflicted mice presented with high amounts of the respective parameters.

### Fluorescence microscopy

Small-intestinal tissue was dissected and luminal contents were flushed out with PBS. Intestines were opened longitudinally and ∼0.5 cm tissue pieces containing macroscopically visible lesions were excised. Tissues were fixed in 1% paraformaldehyde overnight then blocked for two hours at room temperature in blocking solution containing 10% BSA, 1 μg/ml anti-CD16/32, and 0.5% normal rat IgG in PBS. AF647-conjugated anti-Ly-6G antibody (1A8; Biolegend) was added at 0.01 mg/ml in 100 μl PBS per sample, then whole tissue was stained for 24 hours at 4°C. Samples were washed three times with PBS, mounted whole onto slides in Prolong Glass Antifade Mountant (Thermo Fisher Scientific) and cured for two days at room temperature. Images were acquired on a DMI 6000 laser-scanning confocal microscope (Leica) with a 20x NA 0.75 oil-immersion objective. The center of the sample was determined in the Z direction, then imaged. Images were analyzed using ImageJ v2.1.

### RNA sequencing

Intestinal punch biopsies were collected as described above. Five biopsies per sample type were pooled for each mouse. RNA was extracted using the RNeasy Plus Mini Kit (Qiagen). Sequence-ready libraries were prepared using the Illumina TruSeq Stranded Total RNA kit with Ribo-Zero Gold rRNA depletion (Illumina). Quality assessment and quantification of RNA preparations and libraries were carried out using an Agilent 4200 TapeStation and Qubit 3, respectively. Samples were sequenced on an Illumina NextSeq 500 to produce 150–base pair single end reads with a mean sequencing depth of 9 million reads per sample. Raw reads from this study were mapped to the mouse reference transcriptome (Ensembl; Mus musculus GRCm38) using Kallisto v0.46.2^81^. Raw sequence data are available on the Gene Expression Omnibus (GEO; accession no. GSE194334).

All subsequent analyses were carried out using the statistical computing environment R v4.0.3 in RStudio v1.2.5042 and Bioconductor^82^. Briefly, transcript quantification data were summarized to genes using the tximport package^83^ and normalized using the trimmed mean of M values (TMM) method in edgeR^84^. Genes with <1 CPM in 3 samples were filtered out. Normalized filtered data were variance-stabilized using the voom function in limma^85^, and differentially expressed genes were identified with linear modeling using limma (FDR ≤ 0.05; absolute log_2_FC ≥ 1) after correcting for multiple testing using Benjamini-Hochberg.

### Statistics

Statistical analyses were performed using Prism v9.0 (GraphPad Software). Independent groups were compared by Mann-Whitney U test. Survival curves were compared by Mantel-Cox test. Statistical significance is denoted * (p<0.05), ** (p<0.01), *** (p<0.001), **** (p<0.0001), NS (not significant).

## Acknowledgements

We thank Dr. James Bliska for generously providing plasmids for Yop mutant *Yp* strains, as well as Dr. Kimberly Davis for generously providing the mCherry+ *Yp* plasmid. We thank the staff at the PennVet Comparative Pathology Core for their help in preparing the histological samples. We thank Dr. David Christian and Dr. Andrea Stout for key advice on confocal microscopy methods, and Dr. Sunny Shin for constructive editorial comments and scientific discussion. This work was supported by NIH Awards R01AI128530, R01139102A1 (IEB) and a BWF Investigator in the Pathogenesis of Infectious Disease Award (IEB); the Foundation Blanceflor Postdoctoral Scholarship (DS), the Swedish Society for Medical Research postdoctoral fellowship (DS) and the Sweden-America Foundation J. Sigfrid Edström award (DS); NIH NRSA F31AI160741-01 (RM); Mark Foundation Grant 19-011MIA, and F32 AI164655 (JPG); NSF GRFP Award (SP). We thank members of the Brodsky lab for scientific discussion and Dr. Daniel Grubaugh for comments on the manuscript.

## Author contributions

DS established the initial findings of intestinal granulomas. DS, RM, and IEB conceptualized the study and devised experiments. DS and RM devised the methodology and performed experiments. SP, JPG, and IR performed experiments. CAA, ER, and ML performed the histology and histopathological scoring. EK prepared the RNA-sequencing libraries. RM and DB analyzed the RNA-sequencing data. IEB acquired the funding and supervised the study. DS, RM, and IEB wrote the original draft. All authors reviewed and edited the manuscript.

## Competing interests

The author declare no competing interests.

---

**Extended Data Figure 1.**
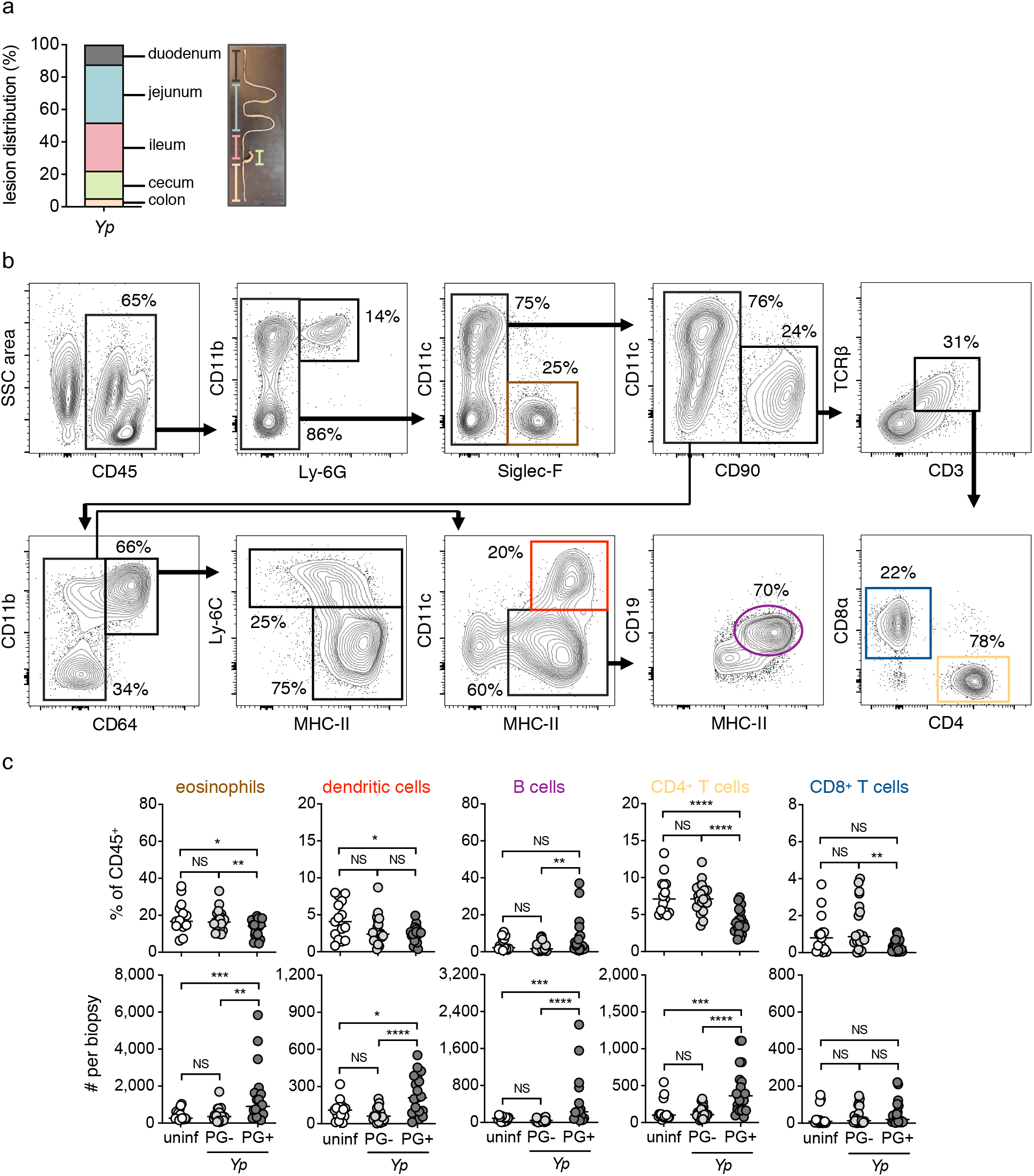
Intestinal pyogranulomas from acutely following oral *Yersinia* infection. (**a**) Frequency distribution (left) of lesions along the intestinal tract, with graphical key of anatomical segments (right). Each colored bar represents the mean frequency of lesions in a given segment (n = 20 mice). Only mice with >9 total pyogranulomas were included. Pooled data from three independent experiments. (**b**) Flow cytometry plots displaying the gating strategy employed to identify eosinophils, dendritic cells, B cells and T cells in small-intestinal tissue. Representative images of four independent experiments. (**c**) Frequency (top) and total number (bottom) of eosinophils, dendritic cells, B cells, CD4^+^ T cells and CD8^+^ T cells (left to right) in small-intestinal uninf, PG- and PG+ tissue. Each circle represents the mean of 3-10 pooled punch biopsies (ø = 2 mm) from one mouse. Lines represent median. Pooled data from four independent experiments. Mann-Whitney U test was performed for all statistical analyses. NS = not significant.

**Extended Data Figure 2.**
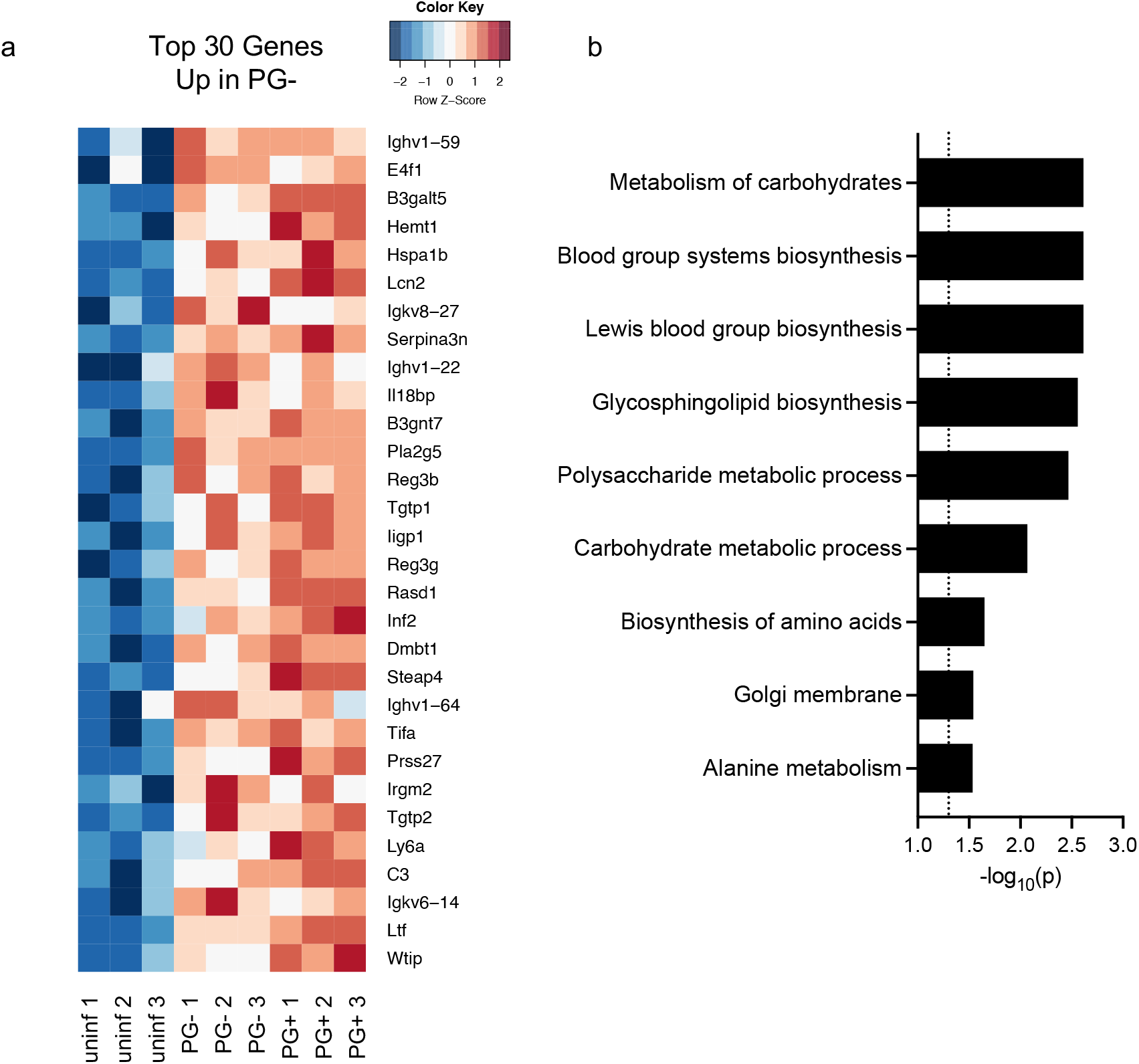
Non-pyogranuloma tissue does not undergo a pro-inflammatory response during intestinal *Yp* infection. (**a**) Heatmap of top 30 significantly upregulated genes in PG- compared to uninfected samples in descending order by fold change. False discovery rate < 0.05 using Benjamini-Hochberg procedure. (**b**) Gene ontology analysis of top 30 upregulated genes by fold change only in PG- compared to uninfected samples. Dotted line denotes p = 0.05.

**Extended Data Figure 3.**
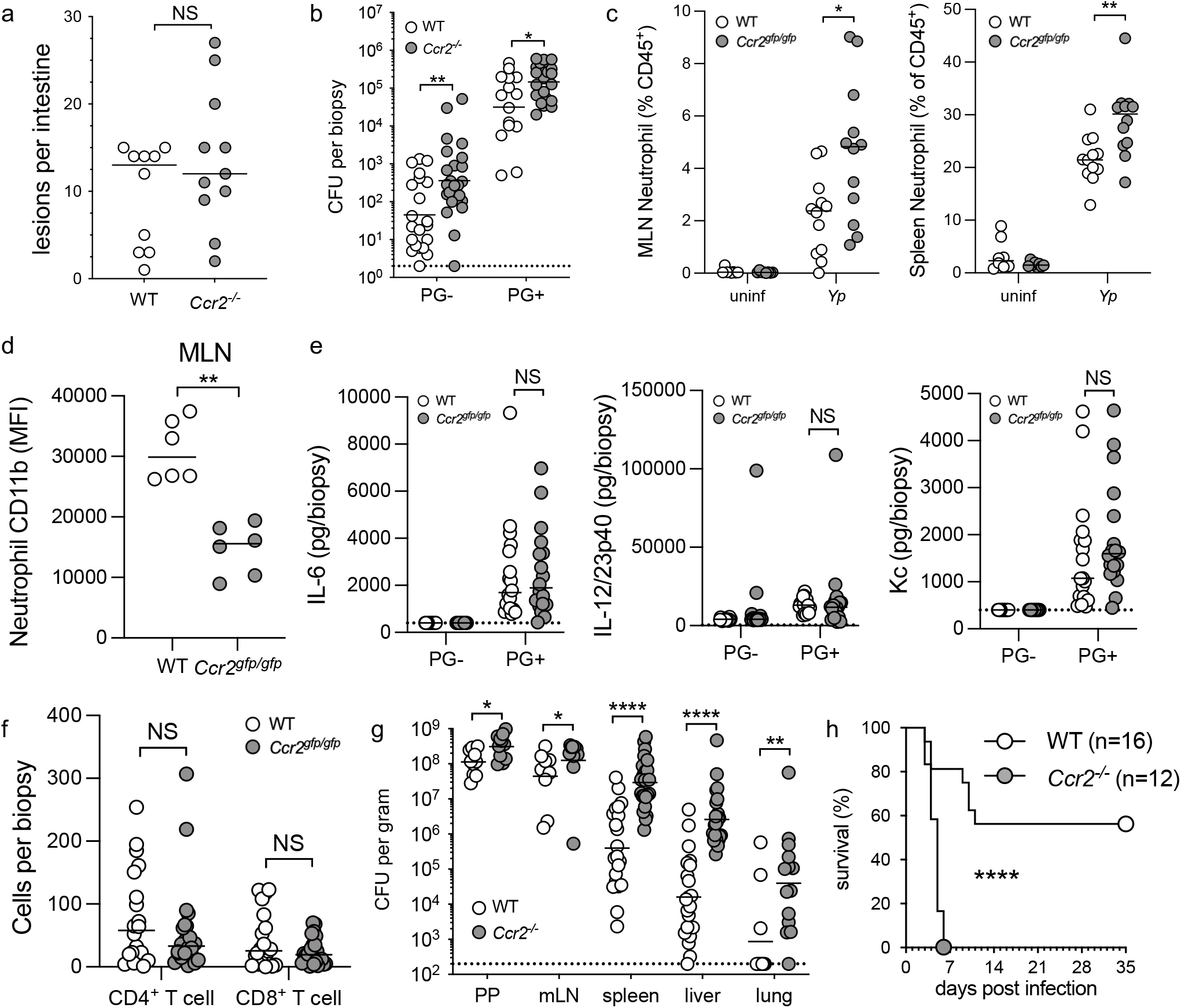
*Ccr2^-/-^* mice have increased susceptibility to oral *Yersinia* infection. (**a**) Quantification of total number of intestinal lesions at day 3 post infection. Each circle represents one mouse. Line represents median. Pooled data from two independent experiments. (**b**) Bacterial burdens in small intestinal PG- and PG+ tissue at day 3 post infection. Each circle represents the mean *Yp*-CFU of 3-5 pooled punch biopsies from one mouse. Lines represent geometric mean. Pooled data from four independent experiments. (**c**) Frequency of neutrophils in MLN and spleen at day 5 post infection. Each circle represents the mean of 3-10 pooled punch biopsies from one mouse. Lines represent median. Pooled data from two independent experiments. (**d**) MLN neutrophil CD11b expression was measured by flow cytometry. Lines represent median. Representative of two independent experiments. (**e**) Cytokine levels in homogenates of tissue punch biopsies at day 5 post infection were measured by cytometric bead array. Lines represent median. Pooled data from three independent experiments. (**f**) Total number of CD4^+^ and CD8^+^ T cells in small intestinal PG+ tissue. Each circle represents the mean of 3-10 pooled punch biopsies from one mouse. Lines represent median. Pooled data from 4 independent experiments. (**g**) Bacterial burdens in indicated organs at day 3 post infection. Each circle represents one mouse. Lines represent geometric mean. Pooled data from four independent experiments. (**h**) Survival of infected mice. Pooled data from two independent experiments. Statistical analyses by (a-g) Mann-Whitney U test or (h) Mantel-Cox test. NS = not significant.

**Extended Data Figure 4.**
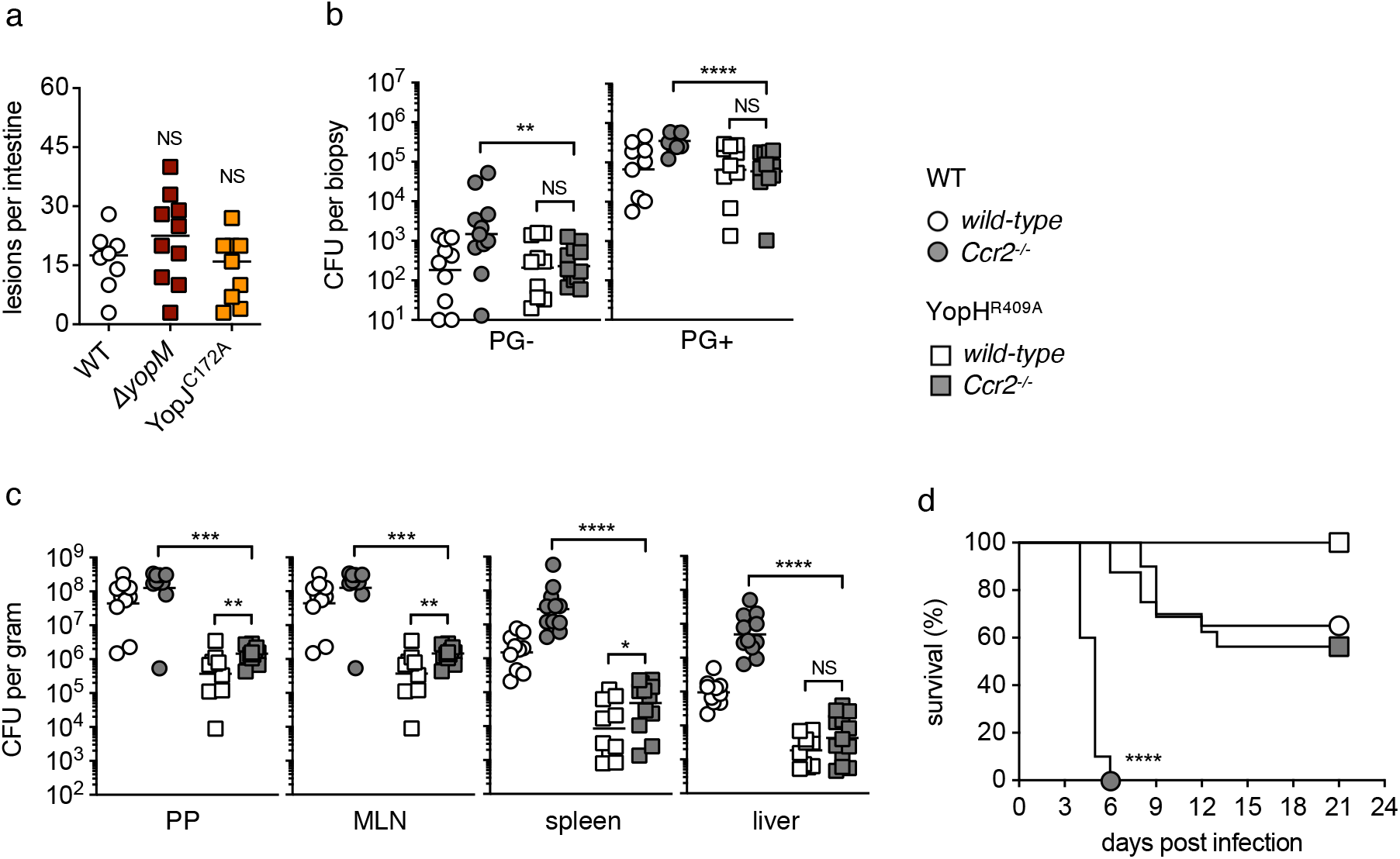
Monocytes are dispensable for restriction of YopH mutant *Yp*. (**a**) Quantification of total number of intestinal lesions upon infection. Each symbol represents one mouse. Lines represent median. Pooled data from 2 independent experiments. (**b**) Bacterial burdens in small-intestinal PG- and PG+ tissue at day 3 post infection. Each symbol represents one mouse. Lines represent geometric mean. Pooled data from two independent experiments. (**c**) Bacterial burdens in indicated organs at day 3 post infection. Each symbol represents one mouse. Lines represent geometric mean. Pooled data from two independent experiments. (**d**) Survival of *Yp*-infected mice. Pooled data from two independent experiments. Statistical analyses by (a-c) Mann-Whitney U test and (d) Mantel-Cox test.

**Extended Data Table 1.** Gene ontology analysis of PG+ vs PG- samples

**Extended Data Table 2.** Gene ontology analysis of PG- vs uninfected samples

**Supplementary code file**

## Notes

### Competing Interest Statement

The authors have declared no competing interest.

### Summary of Updates

Extended data Fig 1a was moved to the main text as Fig. 1h, and text and extended Fig 1 updated accordingly. Additional modifications to introduction text also included to reflect studies on Yersinia pyogranuloma formation.

